# A predicted cancer dependency map for paralog pairs

**DOI:** 10.64898/2026.01.19.700065

**Authors:** Narod Kebabci, Hamda Ajmal, David J. Adams, Colm J. Ryan

## Abstract

**Background:** Genome-wide CRISPR screening has enabled the development of dependency maps in hundreds of cancer cell lines, facilitating the identification of genetic vulnerabilities associated with specific biomarkers. Paralogs, despite being common drug targets, are often missed in these screens as their individual disruption rarely causes a significant fitness defect. Combinatorial screens have revealed that paralog pairs are often synthetic lethal but that these effects are highly context specific. To develop paralogs as therapeutic targets we must identify which paralog pairs are synthetic lethal in which cancer contexts.

**Results:** We develop a machine learning classifier to predict cell-line specific synthetic lethality between paralog pairs. We demonstrate the utility of features derived from the cell-line specific expression and essentiality of the pair and their protein-protein interaction partners for this purpose. We evaluate our predictions across multiple scenarios: predicting for the same pairs in unseen cell lines, for new gene pairs in seen cell lines, and for entirely uncharacterized pairs in unseen cell lines. We show that we can make predictions across all scenarios. We validate our predictions using independent combinatorial CRISPR screens and show that the agreement between our predictions and published experiments approaches the agreement across experiments.

**Conclusions:** Our classifier predicts cell-line-specific synthetic lethality between paralog pairs and provides insights into the underlying features driving these interactions. We make our predictions for 1,005 cell lines available as a resource to facilitate the discovery of context-specific paralog synthetic lethalities and to guide the design of more targeted combinatorial screens.

## Background

Identifying cancer-selective genetic dependencies is a central objective in precision oncology and is key to the development of new targeted therapies [1,2]. Systematic efforts such as the Cancer Dependency Map (DepMap) have used genome-wide CRISPR screening to catalogue the genetic dependencies of over 1,000 molecularly characterised cancer cell lines[3–5]. These efforts have identified new therapeutic targets in specific cancer subtypes and revealed how genetic background and molecular context shape patterns of gene essentiality[4,6,7]. However, many potential therapeutic targets remain undetected in these single-gene knockout screens because the loss of a single gene can often be tolerated due to compensatory relationships with other genes. This is particularly true for paralogs – genes that arose from a gene duplication event[8,9]. Because they arose from a common ancestral gene, pairs of paralogs frequently perform similar molecular functions and hence can compensate for each other’s loss. This can mask cancer vulnerabilities in standard single-gene knockout screens: perturbing one paralog alone may not impair cell viability, even when the combined activity of both is required to support essential processes. This is a significant blind spot in single-gene screens as the majority (>60%) of human protein-coding genes are paralogs as are the majority of targets of approved drugs[10].

Recently, combinatorial CRISPR screens have been employed to simultaneously disrupt hundreds to thousands of paralog pairs across multiple cell lines [9,11–18]. These screens have revealed that many paralog pairs are synthetic lethal – i.e. each gene in the pair can be disrupted individually with relatively little consequence, but their combined disruption causes a severe fitness defect or cell death. Perhaps surprisingly, these synthetic lethal relationships are highly context-specific – often a paralog pair will be synthetic lethal in one cell line but not another. Examples of context-specific synthetic lethalities associated with both cancer-type and mutational status have been reported in the literature[12,14,16,19–21].

Because paralogs are structurally similar, it is likely that many paralog pairs can be co-inhibited using a single small molecule[22,23]. This is the basis of many successful targeted therapies in cancer – such as lapatinib, which inhibits the paralog pair EGFR and ERBB2 and is approved for use in *HER2* amplified breast cancers[24,25], and the MEK inhibitor trametinib, which inhibits the paralogs MEK1 and MEK2 and is approved for use in *BRAF* mutant melanomas[26,27]. Thus, paralog-pair dependencies represent a class of cancer vulnerabilities that are both context-specific and pharmacologically tractable, yet largely invisible to monogenic screening approaches.

Direct experimental mapping of these extremely valuable interactions at scale remains impractical: testing tens of thousands of paralog pairs across a thousand cancer models would require prohibitively large combinatorial screens. Computational prioritization is therefore essential to systematically uncover paralog-pair dependencies and guide experimental validation. We have previously developed a model to predict robust (context insensitive) synthetic lethality for paralog pairs. This model was trained on pairs that were observed to be synthetic lethal across cell lines from multiple cancer types (*pan-cancer* synthetic lethals) and therefore could make accurate predictions for pairs that were observed to be synthetic lethal across most cell lines. However this model could not make predictions about which cell lines a context-specific pair might be synthetic lethal in [28]. Other work has developed models to predict synthetic lethality between non-paralogous gene pairs but these models also typically ignore biological context and cannot make cell-line-specific predictions [28–33].

Here, we develop a random forest classifier to predict cell-line-specific synthetic lethality between paralog pairs. Our model integrates genomic, transcriptomic, and network-based features to capture the cellular context under which synthetic lethal interactions are likely to occur. Using data from existing combinatorial CRISPR screens, we show that our classifier accurately predicts context-dependent synthetic lethality and identifies key features driving these effects. We then apply the classifier to systematically predict the likelihood of synthetic lethality for ∼33,600 paralog pairs across 1,005 cancer cell lines. This large-scale prediction effort provides a comprehensive resource to guide future combinatorial screens, identify tumour-specific vulnerabilities, and support the design of targeted therapeutic strategies based on paralog synthetic lethality.

## Results

### Building a Context-Specific Random Forest Classifier

To develop a machine learning classifier capable of making cell-line-specific predictions of synthetic lethality, we obtained scores for 4,170 paralog pairs in 10 cell lines from a published combinatorial CRISPR screen [12]. We used this to annotate pairs of genes as being synthetic lethal in a specific cell line – i.e. triplets of gene A1, gene A2, and cell line X were labelled as synthetic lethal (SL) or not synthetic lethal (nonSL) (Figure 1). The same gene pair (A1, A2) could be labelled as SL in one cell line and nonSL in another, and indeed this was relatively common. Of the 4,170 paralog pairs, 502 (12.0%) were labelled as SL in at least one cell line and nonSL in another, highlighting the importance of modeling cell-line-specific SL relationships. Notably, *MAP2K1_MAP2K2* was the only paralog pair identified as synthetic lethal across all cell lines. Our predictive features were a combination of cell-line specific features (e.g. relating to the expression of the paralogs themselves in that cell line) as well as cell line agnostic features (e.g. the sequence identity between the two paralogs, which is invariant across cell lines). For cell-line specific features, we made use of the DepMap project [3,34], which provides molecular and phenotypic profiling for hundreds of cancer cell lines. Notably, this project provides genome-wide single-gene CRISPR screen data for most widely used cancer cell lines, which we use to derive input features for our classifier. In effect, we aim to develop a classifier that can answer the following question: Would a paralog pair with these features exhibit synthetic lethality in this particular cell line? In previous work [28], we developed a classifier to predict *robust* synthetic lethality between paralog pairs (i.e. SL that operates across many cell lines from different cancer types), while here we are interested in cell-line specific predictions. We attempted to build our classifier using two distinct strategies – contextualised prediction, where we took as input the predictions output from our previous cell line agnostic classifier and integrated them with cell-line specific information, and full model prediction, where we made use of all of the cell-line agnostic features used to train our previous classifier as well as cell-line specific features (Figure 1).

**Figure 1.**
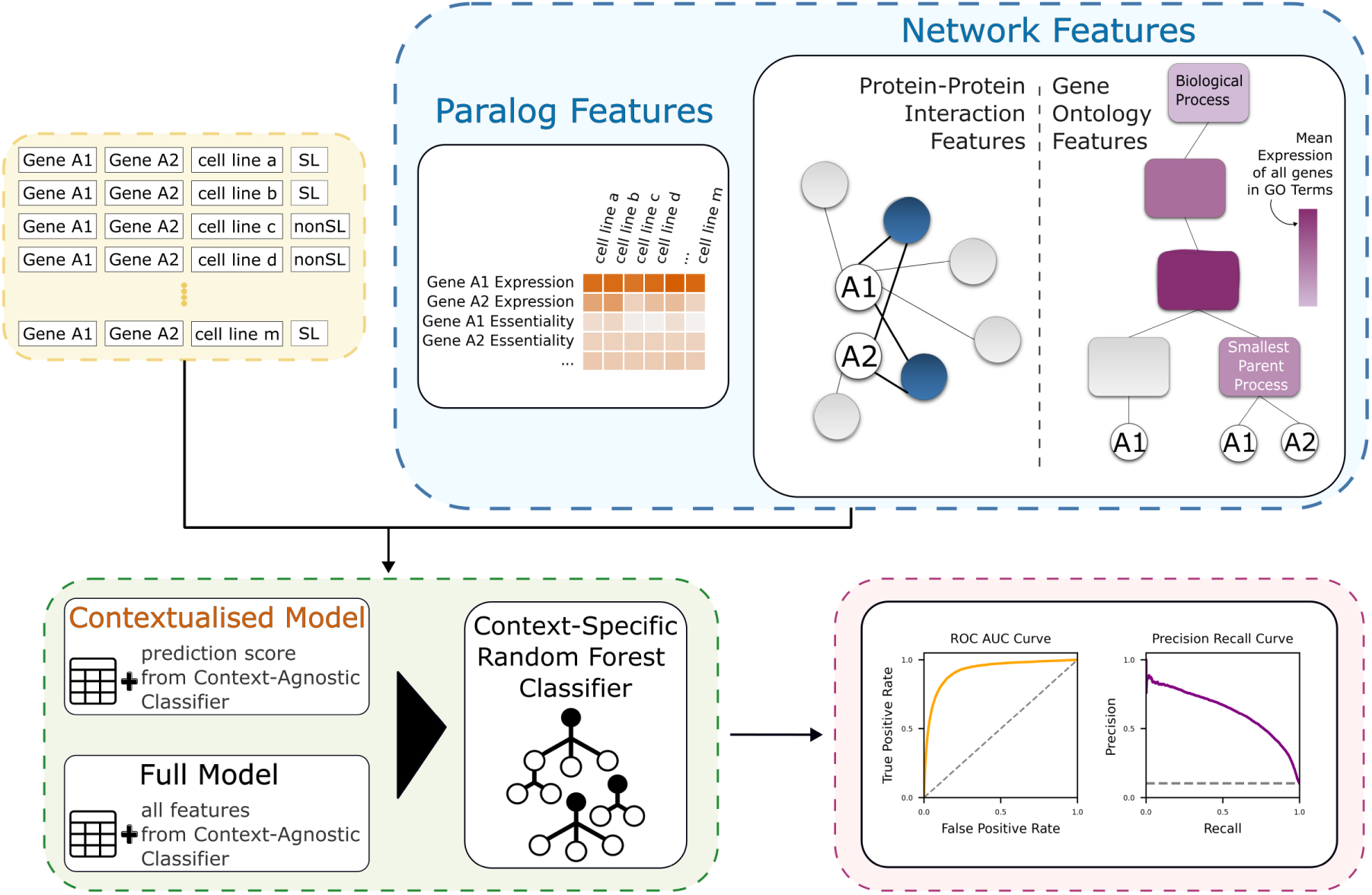
Overview of the context-specific classifier for predicting cell-line specific synthetic lethality between paralog pairs. The input data (yellow box) represents the data obtained from combinatorial CRISPR screens – gene pairs, cell lines, and their corresponding synthetic lethality (SL) classifications derived from the experiments. Feature extraction (depicted in the blue box) encompasses both features of the paralogs themselves, such as their gene expression and gene essentiality, as well as network features derived from protein-protein interaction (PPI) networks and Gene Ontology (GO)-based annotations. Two integration strategies are compared (shown in the green box): contextualised prediction model, which uses prediction scores from a previously published context-agnostic classifier [28], and the full model, which incorporates all context-agnostic features used to train the context-agnostic classifier. A context-specific Random Forest classifier is then trained using these features. The performance of the model is assessed (pink box) using ROC AUC and Precision-Recall curves to evaluate classification accuracy.

### Paralog expression and essentiality are predictive of cell-line specific synthetic lethality

We first assessed whether features relating to the paralogs themselves (such as their mRNA expression level and essentiality in a given cell line) were predictive of cell-line specific synthetic lethality. We evaluated the predictive power of each feature separately, calculating both the Receiver Operating Characteristic (ROC) AUC and average precision for each one (Figure 2a). For features with values assigned to both genes in a paralog pair (A1, A2), we labeled the smallest value as “min” and the largest value as “max”. As a reference, we included sequence identity, a cell line agnostic feature which we previously found to be the most predictive individual feature of robust (non cell-line specific) synthetic lethality [28].

This analysis revealed gene essentiality (min) and gene expression (min) as the most reliable individual predictors of synthetic lethality, achieving ROC AUC values of 0.79 and 0.71, respectively (Figure 2a). Their average precision values ranged from 11% to 7%, reflecting a modest yet meaningful predictive power for SL pairs. It is important to note that the baseline average precision by chance is only 2% due to the class imbalance, as even paralog pairs are rarely synthetic lethal. This means that even well-performing classifiers are expected to have low average precision values in absolute terms. Thus, the observed average precision values for gene essentiality and gene expression are notable and indicate meaningful performance.

The predictive features also showed statistically significant differences in distribution between SL and nonSL pairs based on the Mann-Whitney U test. Our analysis revealed that gene essentiality showed the most significant difference, with SL pairs exhibiting notably lower fitness values than nonSL pairs (*p*=5.9×10^-211^) (Figure 2d). We note that by default, the GEMINI genetic interaction scoring approach used in our training dataset filters out scores for gene pairs where one of the genes causes a significant growth defect when perturbed individually [35]. As a result, the pairs used in our model do not include genes that are individually essential. The ‘gene essentiality’ feature, therefore, suggests that paralog pairs, where at least one member of the pair causes a moderate fitness defect when disrupted individually, are more likely to be synthetic lethal. This finding is consistent with previous work in yeast, where the fitness defect caused by individual gene deletions was associated with genetic interaction degree [36]. Additionally, SL pairs exhibited significantly higher gene expression levels compared to nonSL pairs (*p*=2.3×10^-97^) (Figure 2b). These results underscore the importance of gene essentiality and gene expression as core features in predicting synthetic lethality.

**Figure 2.**
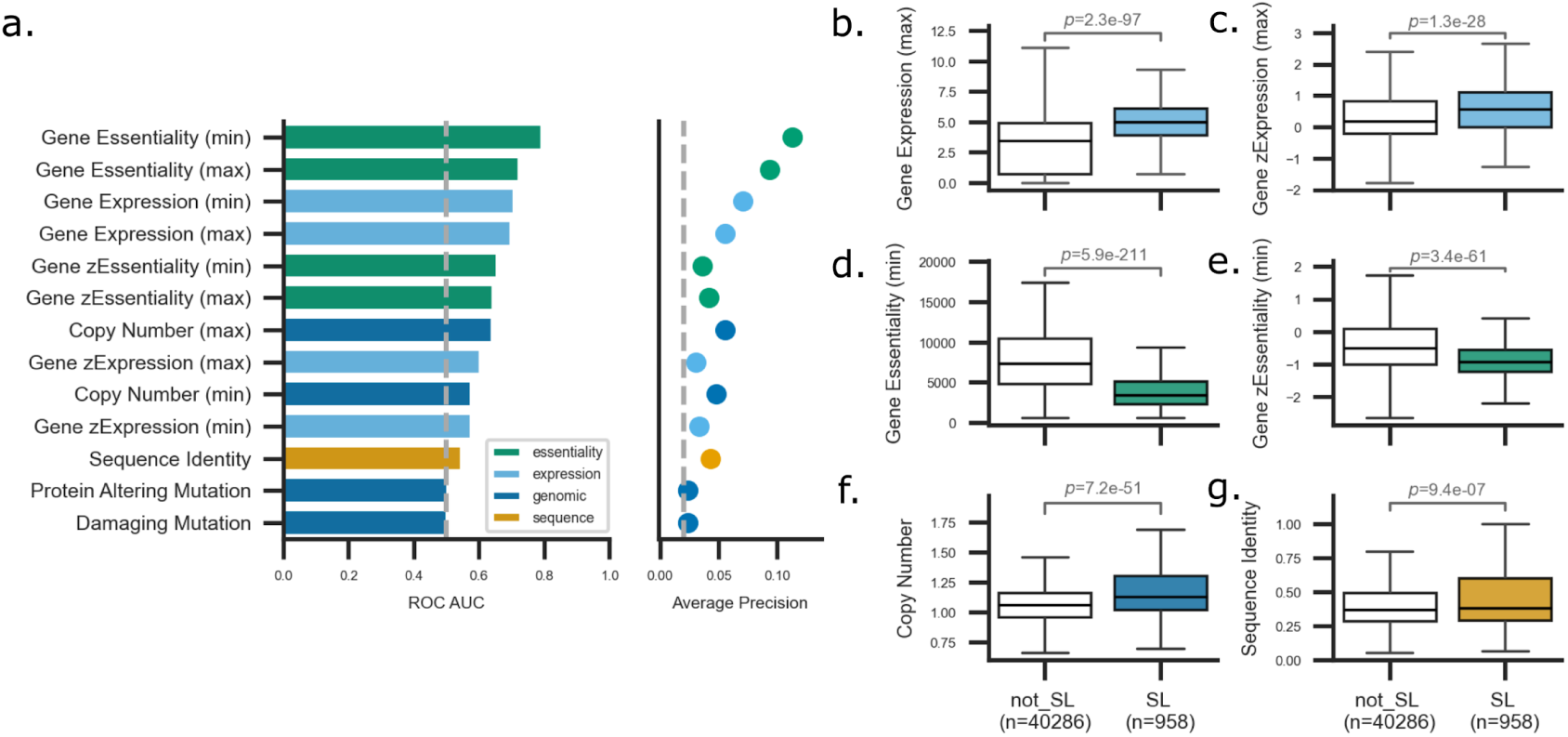
Paralog expression and essentiality are predictive of cell-line specific synthetic lethality. (a) Bar plot showing the performance of individual features measured by ROC AUC scores, and the point plot shows the corresponding Average Precision (AP) scores, evaluated across 4,170 paralog pairs in 10 cell lines. (b-g) Box plots comparing distributions of selected top features between synthetic lethal (SL) and non-synthetic lethal (nonSL) paralog pairs: (b) maximum gene expression, (c) maximum normalized gene expression, (d) minimum gene essentiality, (e) minimum normalized gene essentiality, (f) minimum copy number, and (g) sequence identity. P-values were determined using two-sided Mann-Whitney U tests

For gene expression, we assessed both the measured transcriptomic abundance (TPM) and a normalized version (zExpression), which reflects transcriptomic abundance in each cell line relative to all other profiled cell lines. We found that both were predictive, suggesting it is not just the case that being highly expressed is predictive of SL but also that being *more* highly expressed in the given cell line is predictive. A similar trend was observed for essentiality, where relative essentiality (zEssentiality) was also predictive of SL. These observations were also supported by significant differences between SL and nonSL distributions (zExpression=1.3×10^-28^ and zEssentiality=3.4×10^-61^) (Figure 2c, and Figure 2e, respectively).

While sequence identity emerged as the best standalone predictive feature for robust synthetic lethality, it demonstrates a moderate predictive ability for cell-line specific SL and falls behind cell-line specific features such as gene essentiality and gene expression (*p*=9.4×10^-7^) (Figure 2g). Copy number features show modest performance and limited predictive ability compared to gene essentiality and gene expression, with SL pairs having slightly higher copy number values (*p*=7.2×10^-51^) (Figure 2f). Both protein altering and damaging mutations have near-random performance and negligible average precision, suggesting that mutations in paralogs are not strongly predictive of synthetic lethality in this dataset.

### Network features can distinguish paralog SL pairs from nonSL pairs

Genes, or more precisely, proteins that function together within the same pathway or protein complex, tend to display highly similar genetic interaction profiles, making network-based features valuable for predicting synthetic lethality [37]. We therefore extended our analysis to include network-based features beyond the paralogs themselves, derived from shared protein-protein interactions (PPIs) and shared Gene Ontology (GO) terms. These features generally report on the average essentiality or expression level of the interaction partners of the paralog pair (PPI features) or genes belonging to the same GO term as the pair (GO features).

Among the network-based features, the weighted average gene essentiality of shared protein-protein interactions (PPIs) proved to be the most predictive individual network feature, reaching a ROC AUC of approximately 0.66 (Figure 3a). This feature outperformed its unweighted equivalent from the BioGRID dataset (essentiality of shared PPIs (BG_ALL)), demonstrating the importance of incorporating interaction scores as weights. Similarly, the weighted average gene expression of shared PPIs exhibited strong predictive performance, with a ROC AUC of around 0.65 (Figure 3a). Supporting these performance metrics, we observed that SL pairs had significantly higher values than nonSL pairs for both essentiality (*p*=3×10^^-69^) and expression (*p*=1.9×10^^-62^) of shared weighted PPIs (Figure 3c, and Figure 3h, respectively).

Our approach to gene ontology-based features was inspired by the concept of GO activity scores introduced by [38,39], who proposed that the average expression levels of genes within a biological process can reflect its functional activity across various tissues and cell types. In our research, we adapted this idea by computing both the average essentiality and expression in each cell line for each GO term associated with a paralog pair, with a particular focus on smaller and more specific GO terms. We found that the smallest Gene Ontology Biological Process (GO BP) essentiality and the smallest Gene Ontology Cellular Component (GO CC) essentiality, achieved ROC AUC values of approximately 0.60 and 0.56, respectively (Figure 3a). This was reflected by statistically significant differences between the distributions of scores for these features – smallest GO BP essentiality (Figure 3d) with p-value of 9.0×10^-44 and the smallest GO CC essentiality (Figure 3e) with a p-value of 9.4×10^-24. These features highlight the importance of essentiality within functional pathways and cellular contexts when predicting synthetic lethality.

The smallest GO BP expression values achieved a strong predictive performance, with a ROC AUC of ∼0.60, while the smallest GO CC expression levels exhibited moderate predictive power, with a ROC AUC of around 0.56. Both the smallest GO BP expression (*p*=1.4×10^-29) and GO CC expression (*p*=7.5×10^-23) were also significantly elevated in SL pairs when compared to nonSL pairs (Figure 3f and Figure 3g, respectively). This suggests that interactions at the pathway level may provide more informative signals for synthetic lethality than those from cellular components.

Our findings indicate that paralogs are more likely to be SL in contexts where their protein-protein interaction partners are essential, or where the pathway they function in is essential. They are also more likely to be SL when these functionally related genes are highly expressed.

To assess whether the predictive features are specific to the training dataset or generalizable across different screening experiments, we next examined the predictive performance of the features in an independent combinatorial CRISPR screen. Specifically, we asked whether the same features that performed well in the training dataset [12] also ranked highly in a separate screen from [15], using consistent SL labeling across both datasets (see Methods). This comparison allowed us to assess the robustness of feature-level predictive power across distinct experimental conditions.

We observed strong correlations in ROC AUC values of individual features between the two datasets. Pearson correlation was 0.70 (*p*=0.0038) and a Spearman rank correlation was 0.79 (*p*=0.00047), indicating that many of the same features consistently distinguished SL from nonSL pairs across datasets (Figure S1). These results suggest that features predictive of synthetic lethality in the training dataset, such as gene essentiality and weighted average gene essentiality of shared protein-protein interactions, are reproducible and not limited to a single experimental context.

**Figure 3.**
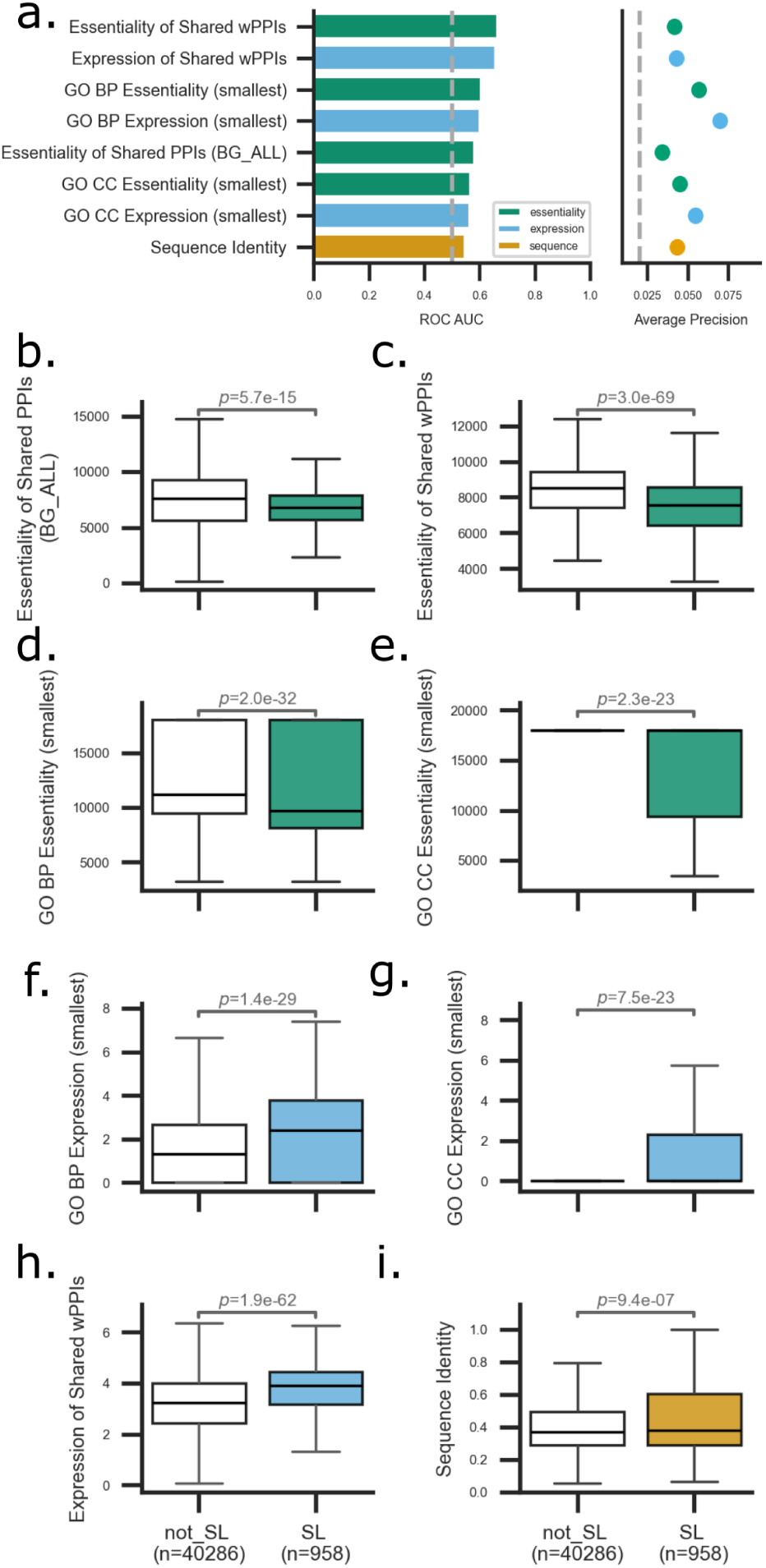
Network-based features from essentiality and expression predict cell-line specific synthetic lethality. (a) Bar plot showing the performance of individual network-based features measured by ROC AUC scores, and the point plot shows the corresponding Average Precision (AP) scores, evaluated across 4,170 paralog pairs in 10 cell lines. (b-i) Box plots compare the distributions of key network-based features between synthetic lethal (SL) and non-synthetic lethal (nonSL) paralog pairs: (b) essentiality of shared PPIs (BG_ALL), (c) essentiality of shared weighted PPIs, (d) smallest GO BP essentiality, (e) smallest GO CC essentiality, (f) smallest GO BP expression, (g) smallest GO CC expression, (h) expression of shared weighted PPIs, and (i) sequence identity. P-values were determined using two-sided Mann-Whitney U tests.

### An integrated classifier predicts cell-line specific synthetic lethality

Having identified individual features capable of predicting cell-line specific SL to varying degrees, we next integrated all features into a random forest (RF) classifier. Four different validation strategies were employed to evaluate the performance of this classifier: (1) randomly selected gene pair/cell line combinations typical of standard machine learning evaluation (Strategy I, Figure 4a) (2) classifier tested with unseen cell lines (strategy II, Figure 4b) (3) classifier tested with unseen paralog pairs (strategy III, Figure 4c); and (4) classifier tested with both unseen cell lines and paralog pairs (strategy IV, Figure 4d). Strategy I reflects a standard machine learning approach where data is simply withheld at random. Strategy II reflects the scenario where a set of paralog pairs has been screened in a subset of cell lines, and the goal is to make predictions for these same paralog pairs in new, unscreened cell lines. Strategy III reflects the scenario where a subset of paralog pairs have been screened in a set of cell lines, and the goal is to make predictions for unscreened paralog pairs in the same set of cell lines. Strategy IV is perhaps the most important assessment, as it aims to make predictions for paralog pairs that have yet to be screened in any cell line, using cell lines that have yet to be screened for any paralog pairs.

The features used for training the *contextualised prediction* models included gene expression, gene essentiality, copy number variations, mutation profiles, sequence identity, and network-based characteristics, alongside the published prediction score from the context-agnostic classifier[28]. In effect what we are trying to do is to take the predictions from the context-agnostic classifier and combine them with cell-line specific information to make them more context-specific. In contrast, the *full prediction* model included all features used to train the context-agnostic classifier, in addition to the features used in the contextualised model, allowing for a more complete integration of all predictive features. The primary goal of both approaches was to predict the probability that a target paralog pair would be synthetic lethal or not within a specific cancer cell line.

Both the contextualised and full model RF classifiers achieved a high ROC AUC of 0.92, 0.93, and an average precision of 36%, 41% respectively (Strategy I, Figure 4a), outperforming individual features when using random splits of paralog pairs and cell lines. In comparison, the context agnostic classifier performed less effectively, recording an ROC AUC of 0.68. Although sequence identity was previously identified as highly predictive of robust synthetic lethality, it proved to be a relatively poor predictor (ROC AUC of 0.54 and an average precision of ∼4%). This highlights the importance of context-specific biological data for achieving accurate predictions.

The classifiers maintained high performance even in challenging scenarios, such as Strategies II, III, and IV. Across all validation scenarios, the context-specific RF classifiers (contextualised and full) consistently outperformed the context-agnostic classifier and sequence identity, confirming the importance of integrating context-specific biological data.

**Figure 4.**
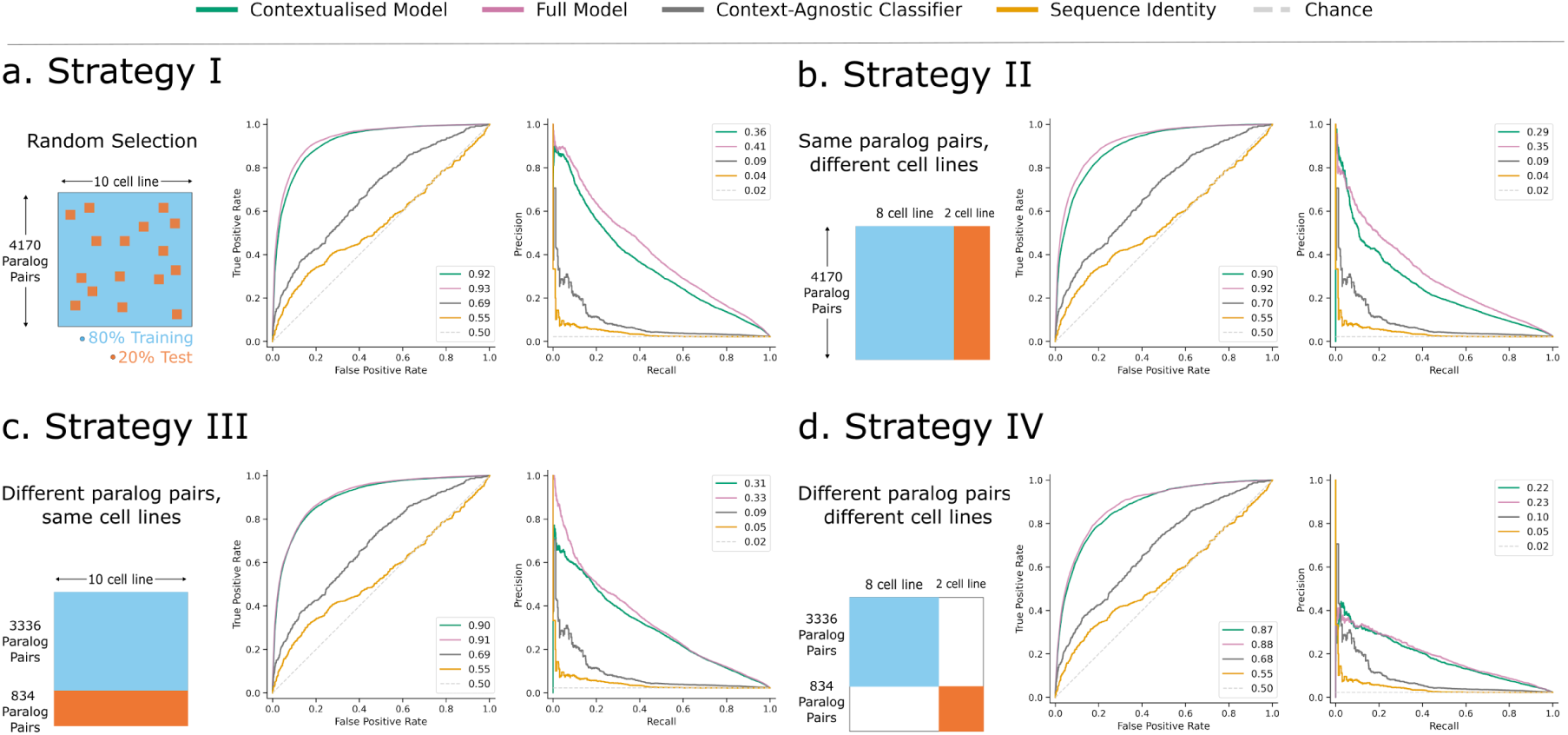
Performance evaluation of the context-specific random forest classifiers across four validation strategies. ROC and Precision-Recall curves are shown for four models: (a) Strategy I, random selection of gene pairs and cell line combinations (80% training, 20% test), (b) Strategy II, seen paralog pairs evaluated in unseen cell lines, (c) Strategy III, unseen paralog pairs evaluated in previously seen cell lines, (d) Strategy IV, unseen paralog pairs evaluated in unseen cell lines. Classifiers include the contextualised model (green), full model (pink), context-agnostic classifier (gray), and sequence identity (orange). The dashed line indicates random performance (chance).

In terms of ROC AUC, the best performance was observed for Strategy I (AUC = 0.92) with the lowest performance observed for Strategy IV (AUC = 0.87), and Strategy I and II falling in between (AUC = 0.9 for both). To assess whether these differences were statistically significant, we evaluated ROC AUC and average precision scores over multiple iterations of both contextualised and full models. The distribution of these scores for each evaluation strategy is visualized as box plots in Figure S2. This revealed a significant difference between Strategy I and both Strategy II and III, no significant difference between Strategies II and III, and a significant difference between Strategy IV and both Strategies II and III (Figure S2a). This suggests that traditional machine learning evaluation, where data points are removed and random, may overestimate performance. Moreover, predicting for unseen paralog pairs in unseen cell lines is more challenging than predicting in either scenario alone. Nonetheless, the ROC AUC is high across all scenarios, ranging from 0.92 in Strategy I to 0.87 in Strategy IV.

A similar pattern was observed for average precision values (Figure S2b). The performance declined significantly from Strategy I to Strategy II (p = 3.3×10^-10). The difference between Strategy II and Strategy III was not statistically significant (p = 0.051), mirroring the ROC AUC results. However, the drop from Strategy III to Strategy IV was again statistically significant (p = 3.9×10^-9), underscoring the compounded difficulty of dual generalization.

These findings confirm that while the classifiers maintain strong predictive performance in more restricted validation scenarios, their accuracy diminishes slightly under increasingly stringent evaluation conditions. The comparable performance of Strategy II and Strategy III suggests that generalizing to either unseen cell lines or unseen paralog pairs presents a similar level of difficulty. However, predicting entirely new cell line-pair combinations, as in Strategy IV, remains the most challenging task. By default, Strategy IV is trained with less data than the other models (because both cell lines and paralog pairs are excluded), suggesting that the poorer performance may simply reflect the smaller training data. However, after artificially reducing the training data for the other models, Strategy IV still performs worse overall, suggesting that predicting on the combination of new cell lines and new pairs is more challenging (Figure S3).

### Evaluating the Performance of the Context-specific Classifier on an Independent Dataset

To maximize the model’s learning potential, we trained the context-specific RF classifier using our complete primary dataset without internal partitioning. We then assessed the performance of our context-specific RF classifier using an independent combinatorial CRISPR screen from Klingbeil *et al* [18]. From this screen, we created two distinct test sets, each consisting of data from 20 cell lines that were not used during the training process. We excluded one overlapping cell line (A549, DepMap ID: ACH-000681), which was part of the training data, as well as another cell line (NCIH1436, DepMap ID: ACH-000830), due to missing gene essentiality data. The seen set included 692 paralog pairs that were also present in the training data, while the unseen set contained 1,012 paralog pairs that were entirely novel for the classifier. This setup enabled us to evaluate the generalization ability of the classifier across both seen and unseen paralog pairs across entirely unseen cell lines.

**Figure 5.**
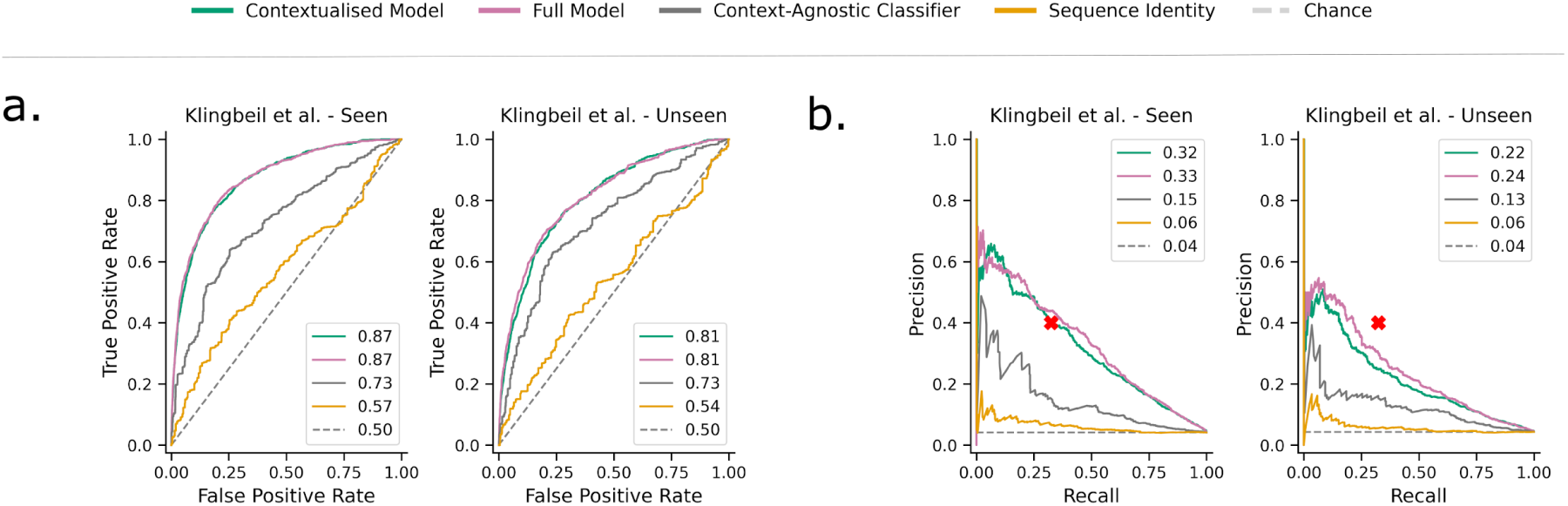
Context-specific classifiers outperform baseline models on independent data, including unseen paralog pair–cell line combinations. (a) ROC curves and (b) precision-recall curves show the performance of contextualised and full model classifiers on the Klingbeil et al. (2024) dataset. Both classifiers outperformed the context-agnostic classifier and the sequence identity baseline across test sets of *seen* and *unseen* paralog pairs, all evaluated on cell lines not used during training. The red cross on the precision-recall plot providing reference for the maximum reproducibility expected from experimental agreement between screens.

We compared the performance of the context-specific RF classifiers (both contextualised and full models) against two baselines: sequence identity and the prediction scores calculated by the context-agnostic classifier. As shown in Figure 5, the context-specific classifiers consistently outperformed the baseline classifiers across both the seen and unseen subsets of the Klingbeil study.

On the seen subset of the Klingbeil study, the context-specific classifiers achieved ROC AUC scores of 0.87 (contextualised) and 0.87 (full), and average precision of 32% (contextualised) and 33% (full), respectively. In contrast, the context agnostic classifier achieved a ROC AUC of 0.73 and an average precision of 15%, while sequence identity achieved only a ROC AUC of 0.57 and an average precision of 6%.

On the unseen subset of the Klingbeil study, although performance dropped, as expected in this more challenging generalization task, the context-specific classifiers still retained meaningful predictive power with ROC AUC scores of 0.81 (Figure 5a) for both contextualised and full models, and average precision of 22% (contextualised) and 24% (full). The context-agnostic classifier again achieved lower performance with an ROC AUC of 0.73 and an average precision of 13%. Sequence identity performed poorly compared to others, with an ROC AUC of 0.54 and an average precision of 6%. These results demonstrate that incorporating cell-line-specific information provides a consistent advantage, even in completely novel contexts.

To place these results in a biological context, we calculated the empirical agreement between the training and test datasets, specifically examining how often synthetic lethal interactions were reproducibly identified in both sets when the same cell line was screened. This overlap resulted in a precision of 0.40 and a recall of 0.32. These values demonstrate the reproducibility of the findings across experiments and serve as a reference point for evaluating classifier performance (red cross in Figure 5b). As in Figure 5b, our predictions for previously seen paralog pairs in new cell lines achieve the same level of precision and recall, while those for previously unseen paralog pairs in new cell lines fall just below this.

We further tested the generalizability of our classifier using an additional independent screen [15] which screened ∼1,000 paralog pairs in two cell lines not included in the training dataset (Figure S4). Again we found that our context-specific classifiers outperformed sequence identity alone and the context-agnostic classifier.

These findings highlight the robustness of the context-specific RF classifiers across diverse datasets while also suggesting room for improvement in its performance on completely unseen data. Refining feature selection, increasing training data, and exploring additional context-specific annotations may enhance its predictive capabilities.

### Associating paralog synthetic lethality with specific biomarkers in breast cancer

Having demonstrated that the concordance between the predictions of our classifier and published experiments is comparable to the agreement across experimental datasets, we next sought to make predictions for 33,574 paralog pairs across all 1,005 cell lines with sufficient data (a total of 33,541,469 predictions). This represents the subset of approximately 36,000 paralog pairs we previously predicted that have the necessary context specific information required for the classifier (i.e. those with gene expression quantification and gene essentiality data). As performance was slightly higher for the full model, we used this model to make all predictions.

To explore the utility of this approach for identifying vulnerabilities associated with cancer subtypes and mutations, we focused on *HER2* amplification in breast cancer. We stratified the breast cancer cell lines according to *HER2* status (see methods), and then sought to identify paralog pairs that were predicted to be synthetic lethal with greater confidence in the *HER2* amplified group. We identified all pairs with a high difference in prediction score between the two groups (difference between the median score in *HER2* amplified models vs median score in WT models > 0.1) and further filtered to those pairs that had a nominally significant association between *HER2* status and prediction score (p<0.05, Mann-Whitney U test). This resulted in 185 pairs whose predicted synthetic lethality appeared dependent on *HER2* status. These include high confidence predictions between *HER2* and its paralogs *EGFR* and *ERBB3*, pairs with well established synergistic relationships [40–43]. Furthermore *AKT1* and *AKT2* are predicted to be synthetic lethal in the HER2 amplified models, consistent with the known role of HER2 in activating AKT signaling [44]. Beyond these established pairs, many other pairs are predicted to be selectively lethal in the *HER2* background, including *AP1G1*_*AP1G2,* and *PTP4A1_PTP4A2*. *AP1G1* encodes the γ1-Adaptin subunit of adaptor protein complex-1, recently shown to be overexpressed in breast tumours, associated with relapse-free survival, and specifically highly expressed in *HER2*+ tumours[45]. *PTP4A1* and *PTP4A2* encode PRL1 and PRL2, phosphatases with an established role in AKT signaling [46] and potentially a direct role in establishing HER2*+* tumours – transgenic mice co-expressing PTP4A2 and ErbB2 showed accelerated mammary tumour formation compared to ErbB2 alone [47]. These make attractive candidates for further exploration of synthetic lethality in HER2-amplified breast cancer cell lines. In contrast to these selective dependencies, other established synthetic lethals are predicted with high-confidence across all cell lines but no significant difference between *HER2* amplified and non amplified – e.g. *ASF1A_ASF1B* and *SAR1A_SAR1B* (which were both identified as context-insensitive synthetic lethals in a recent meta-analysis [17]).

**Figure 6.**
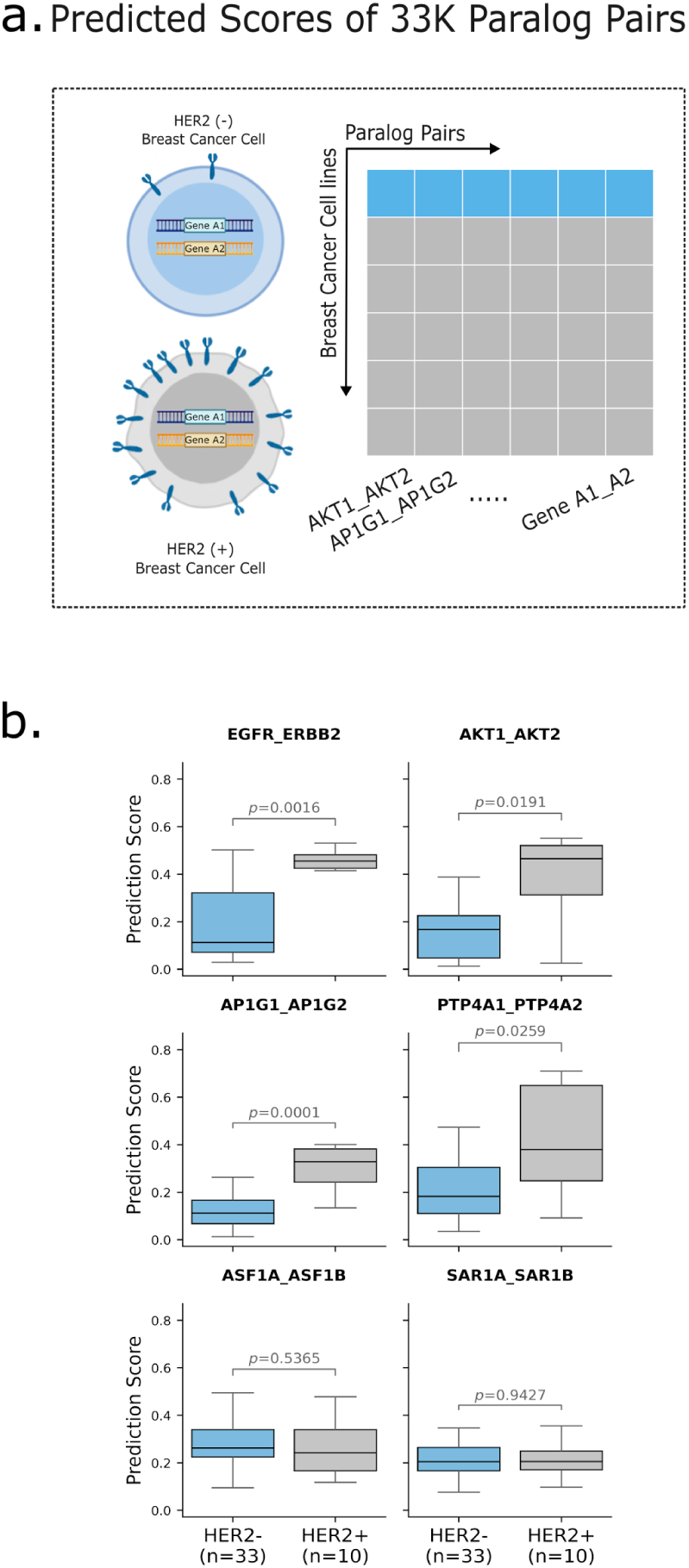
Synthetic lethality predictions reveal HER2-associated paralog dependencies and novel selective interactions in breast cancer. (a) We generated a comprehensive map of predicted synthetic lethality (SL) scores across ∼33,000 paralog pairs and ∼1,000 cancer cell lines. (b) The boxplots show SL prediction scores generated by the full model for selected gene pairs across HER2-positive (n=10) and HER2-negative (n=33) breast cancer cell lines. Known HER2-associated paralog interactions (*EGFR_ERBB2*, *AKT1_AKT2*), novel HER2-selective predictions (*AP1G1_AP1G2*, *PTP4A1_PTP4A2*), and pan-cancer synthetic lethal pairs (*ASF1A_ASF1B*, *SAR1A_SAR1B*) are shown. P-values are from the Mann–Whitney U test.

## Discussion

Understanding the genetic dependencies that underlie cancer cell fitness is essential for developing new targeted therapies. Paralog pairs represent a major blind spot in existing dependency maps derived from monogenic CRISPR screens because functional compensation often masks essential functions that are only revealed when both paralogs are perturbed. Since these paralog pair dependencies are potentially therapeutically tractable, their absence is a significant limitation of existing resources. Here, we develop a cell-line–specific machine learning classifier that predicts the synthetic lethality of paralog pairs by integrating genomic, transcriptomic, and network features. Recent combinatorial CRISPR screens have revealed that these relationships are highly context-specific, and we show that they can be predicted from features that capture the cellular environments in which paralog pairs operate.

Our results highlight several general principles. The expression patterns of the paralogs and their interaction partners are among the strongest predictors, underscoring the role of regulatory context in shaping paralog compensation. Essentiality profiles provide additional information, suggesting that the fitness consequences of disrupting interaction partners and pathway members help predict whether a paralog pair will be synthetic lethal.

We compared two modelling strategies: a contextualised model, which refines predictions from a prior context-agnostic classifier using cell-line-specific features, and a full model, which jointly incorporates all global and context-specific features. The full model slightly, but consistently, outperformed the contextualised model, indicating that contextual information should be integrated directly during training rather than added post hoc. Our finding that prediction is hardest for the combination of unseen paralog pairs in unseen cell lines suggests that as coverage of both increases, so will the potential accuracy of our predictions. Overall, model performance approaches the agreement observed between independent combinatorial CRISPR experiments, suggesting that predictive accuracy is nearing the limits imposed by experimental noise.

Although our study focuses on paralogs, the same modelling framework can be adapted to non-paralog gene pairs, which constitute a much larger space of potential context-specific synthetic lethal interactions. Many of the most informative features in our model—such as expression, essentiality patterns, and protein-interaction context—are likely to be equally relevant for predicting conditional dependencies between any two genes. The main constraint on extending this approach genome-wide is the current lack of large, high-quality combinatorial perturbation datasets for non-paralog pairs. Such datasets are beginning to emerge in individual cell lines[48–50], but as they become available across multiple cell lines, the feature-integration and context-aware learning strategies developed here could be applied to identify context-specific synthetic lethality at scale.

## Conclusions

We developed a context-specific classifier that predicts cell-line-specific synthetic lethality between paralog pairs and provides insights into the underlying features driving these genetic interactions. We make our predictions for 1,005 cell lines publicly available as a resource to facilitate the discovery of context-specific paralog synthetic lethalities and to guide the design of more targeted combinatorial screens.

## Methods

### Data Collection and Labeling

*Combinatorial CRISPR screens.* We collected screen results from three combinatorial CRISPR screens [12,15,18]. [12] and [18] used the GEMINI sensitive scoring method, which we recently demonstrated as performing well in identifying known synthetic lethal pairs from combinatorial screens [51]. The published Parrish screen made use of an alternative scoring system, and so we reprocessed the raw read count data from this screen using the GEMINI sensitive scoring method to make it comparable to the other two screens [35,52].

*Labelling strategy.* Previous work has established that many gene pairs that display a negative genetic interaction score do not actually cause a significant growth defect, typically occurring when the genes under assessment cause a weak fitness increase when perturbed individually and a near neutral effect when perturbed in combination[11,28]. Therefore, to label paralog pairs as synthetic lethal (SL) or non-synthetic lethal (nonSL), we applied a dual threshold approach based on both the GEMINI interaction score and the severity of the growth defect. For each individual study, we derived a list of essential genes by combining the Hart essential gene list [52] with the set of essential gene controls provided by the study authors. To generate a screen specific log fold change (LFC) threshold, we calculated a mid-point between the median LFC values of constructs where a single targeted gene is a known essential and the median LFC values of constructs where neither targeted gene is a known essential. Additionally, for each screen, we calculated a GEMINI threshold based on the 90th percentile of GEMINI scores within that study. Only paralog pairs with GEMINI scores above this threshold and LFC values below the screen specific LFC threshold were labeled as synthetic lethal (SL). All other pairs were labeled as nonSL.

### Preprocessing and Feature Engineering

Genomics data. Gene expression, gene essentiality, copy number, and somatic mutation profiles were obtained from the Cancer Dependency Map (DepMap) 22Q4 release [53]. Gene expression values were extracted from the file OmicsExpressionProteinCodingGenesTPMLogp1.csv, gene essentiality scores were sourced from CRISPRGeneEffect.csv, gene-level copy number data were obtained from OmicsCNGene.csv, and somatic mutation profiles were retrieved from the file OmicsSomaticMutations.csv, which were generated using the Mutect2 pipeline. For each gene in each cell line, mutation status was encoded as two binary features. A gene was labeled mutated and assigned a value of 1 if any variant annotation (e.g., missense, splice site) was reported for that gene in the given cell line, and 0 otherwise. The same gene was labeled as damaging and assigned a value of 1 if any of its variants in that cell line were flagged as CCLEDeleterious or LikelyLoF, and 0 otherwise. These features were not mutually exclusive, as damaging mutations represent a subset of all reported variants.

For cell lines without available gene essentiality data in DepMap, i.e. HeLa and PC9, data were retrieved from [54,55], respectively. To standardize the data, all essentiality scores were ranked, and the ranked scores were used in further analysis. For the scores obtained from the DepMap portal, a lower score indicates a higher essentiality, meaning that as the ranks increase, the essentiality decreases. The same applies to the CRISPR score used for scoring PC9. However, for the HeLa screen, which utilizes Bayes Factors, a higher score indicates greater essentiality. Therefore, the HeLa screen was ranked in reverse order.

*Protein-protein interaction and gene ontology data.* We retrieved protein-protein interactions from the STRING database [56] (full network is downloaded on December 18, 2022, version 11.5). Protein identifiers were mapped to their corresponding Ensembl gene identifiers using data from the HGNC website [57]. We only consider interactions with a confidence score higher than 200. We also downloaded protein-protein interactions from the BioGRID database [58] (all interactions are downloaded on April 29, 2023, version 4.4.221). Gene Ontology (GO) biological process and cellular component data were downloaded from the Ensembl BioMart (downloaded on September 11, 2023).

*Feature values and predictions from the context-agnostic classifier.* We downloaded feature values and prediction scores for ∼36.6k paralog pairs from [28], a random forest classifier developed to predict robust (context-agnostic) synthetic lethality between paralog pairs in cancer. This classifier, referred to throughout this study as the context-agnostic classifier, served as a baseline reference model for comparison.

*Weighted averages of shared protein-protein interactions.* For each paralog pair, we identified common interactors using the weighted STRING protein-protein interaction (PPI) network. The interaction scores were computed as the product of confidence scores between each shared interactor and the individual genes in the paralog pair. For example, if protein A1 interacts with protein B with a confidence score x, and protein A2 interacts with protein B with a confidence score y, the interaction score for protein B as a shared interactor of the paralog pair (A1, A2) is calculated as x*y. Using these interaction scores as weights, we calculated weighted averages for both gene essentiality and gene expression of the common interactors for each paralog pair. This was achieved by multiplying the corresponding values (essentiality or expression) of the common interactors by their interaction scores, summing the resulting weighted values, and dividing by the sum of the interaction scores.

*Averages of the shared Gene Ontology terms.* We identified biological and cellular processes associated with paralog pairs by integrating Gene Ontology (GO) annotations. GO terms were filtered to include only processes with at least 5 but fewer than 100 associated genes. For all GO terms linked to paralog pairs, we calculated the average gene essentiality and expression of genes sharing those GO terms across all cell lines.

Each paralog pair could be annotated to multiple GO terms, and so for each pair in each cell line, we derived three GO-related features based on GO annotations: *GO_smallest*, *GO_max*, and *GO_min*. *GO_smallest* corresponds to the GO term with the smallest gene set size among the terms associated with the paralog pair. *GO_max* represents the GO term associated with the highest average score (either gene essentiality or gene expression). *GO_min* corresponds to the GO term associated with the lowest average score (either gene essentiality or gene expression).

*Labelling HER2 status of Breast cancer cell lines.* We filtered breast cancer cell lines based on their *OncotreeLineage* annotations provided in the DepMap 22Q4 metadata (Model.csv). By using the additional subtype annotations from *LegacySubSubtype,* we generated HER2 labels. Cell lines labeled as ERneg_HER2pos or ERpos_HER2pos were defined as HER2-positive, while those labeled as ERneg_HER2neg or ERpos_HER2neg were defined as HER2-negative. Cell lines with unrecognized subtypes were labeled as ‘unknown’ and excluded from further analysis. This labeling was used to stratify breast cancer cell lines for downstream evaluation.

### Random Forest Configuration

We developed contextualised prediction and full model classifiers using a Random Forest (RF) model, which was implemented using the RandomForestClassifier function from the scikit-learn library (version 1.2.2).

*Hyperparameter Tuning.* We conducted a grid search over a predefined hyperparameter space to determine optimal configurations. Utilizing the GridSearchCV function from the scikit-learn library (version 1.2.2) [59], we focused on optimizing our models based on average precision (AP) as the scoring metric. We conducted hyperparameter tuning separately for all four evaluation approaches (Strategy I to IV), using grid search combined with repeated stratified cross-validation, with model selection based on the mean average precision across cross validation folds. After completing the grid search for each evaluation strategy (I–IV), we performed t-tests on the AP scores to identify hyperparameter configurations whose performance was not significantly different from the best-performing configuration for each model. We then retained only those configurations that were shared across all four models. From this final set, we selected the configuration that achieved the highest performance for all models. We evaluated the following hyperparameters using grid search: n_estimators (600, 800, 1000), max_depth (7, 10, 15, 20), min_samples_leaf (4, 8, 12), and max_features (0.1, 0.2, 0.4). The optimal configuration across the all four models was determined as: n_estimators=600, max_features=0.2, max_depth=20, and min_samples_leaf=4.

### Model Training and Design

We developed two machine learning classifiers, contextualised prediction and full model, to predict context-specific synthetic lethality between paralog pairs. These classifiers differ in their feature engineering approaches, but they are both trained on the same combinatorial CRISPR screen data [12]. Additional file 1 presents a comparative summary of the features used in both models.

The contextualised prediction model used a total of 17 features, including gene expression, gene essentiality, copy number, mutation profiles, sequence identity, the published prediction score from the context-agnostic classifier [28], and six network-based features to train the classifier. In contrast, the full model utilized the complete feature set used to train the context-agnostic classifier, rather than incorporating only the prediction score as a feature, alongside context-specific genomic and network features. Genomic features, including gene expression, essentiality, copy number, and mutation calls, were extracted separately for each member of the paralog pair (A1 and A2). In contrast, network-derived features—such as those based on shared protein-protein interactions and gene ontology annotations—were computed once per pair, representing properties of the gene pair as a whole rather than individual genes.

*Training and Evaluation data.* The screens from [12] were used to build a context-specific random forest (RF) classifier, while other combinatorial CRISPR studies served as independent validation datasets to assess the classifier’s performance. Although [12] screened 5,065 gene pairs across 11 different cell lines, we used 4,170 gene pairs and 10 cell lines to build our classifier, as not all genes had available gene essentiality, expression, or copy number data in the DepMap portal. We excluded the MEWO cell line (DepMap ID: ACH-000987), a melanoma cell line, from both the training and evaluation datasets because of the lack of gene essentiality data in the DepMap portal and other resources. Additionally, we removed gene pairs identified as common essentials in the DepMap portal, as their individual disruption already leads to a growth defect in the cell. Ultimately, the training dataset consisted of 958 data points labeled as SL and 40,286 data points labeled as nonSL.

### Data Splitting and Cross-validation

We used four distinct 5-fold cross-validation strategies, to assess the classifiers’ effectiveness. In each strategy, we varied the grouping criteria used to separate training and test folds. This allowed us to simulate different levels of data novelty.

To ensure a balanced representation of SL and nonSL pairs across the folds, we used StratifiedKFold and StratifiedGroupKFold from the scikit-learn library (v1.2.2). This process was repeated 10 times to reduce variance in performance estimates and obtain a reliable assessment of the classifiers’ effectiveness. The dataset splits for different evaluation strategies were structured as follows:

● Strategy I (Random Partitioning): We applied StratifiedKFold without grouping. This represents a standard random split where distinct data points are tested, but the same cell lines and gene pairs appear in both training and testing sets.
● Strategy II (Unseen Cell Lines): We applied StratifiedGroupKFold grouping by Cell Line ID. In this setup, the test set consists entirely of cell lines never seen by the model during training, though the gene pairs themselves may have been encountered in other cell lines.
● Strategy III (Unseen Gene Pairs): We applied StratifiedGroupKFold grouping by Gene Pair ID. Here, the test set contains only gene pairs that are completely novel to the model, simulating the prediction of new synthetic lethal interactions.
● Strategy IV (Unseen Cell Lines and Unseen Gene Pairs): A custom cross-validation function was developed to ensure that both unseen cell lines and unseen gene pairs were included in the test fold.

Because of the different ways of splitting the data, Strategies I-III contained similar numbers of samples in their train/test splits (∼33,000 points for training and ∼8,200 points for testing) while Strategy IV contained fewer samples (∼26,400 for training and ∼1,650 for testing). For the analysis in Figure S3 we artificially restricted the points for training in Strategies I-III to match the number in Strategy IV.

### Model Evaluation

*Evaluation metrics.* Model performance was assessed using the Receiver Operating Characteristic (ROC) AUC, and Precision-Recall (PR) metrics. For each fold and repetition, ROC AUC and average precision metrics were computed and averaged to characterize the performance of the contextualised and full classification models. Next, the averaged values are utilized to visualize the ROC and PR curves.

*Statistical testing.* To compare the predictive performance of the contextualised and full model classifiers with each other and against the context-agnostic classifier baseline, we used two-sided paired t-tests. These tests were conducted on the ROC AUC and average precision obtained from repeated cross-validation runs to assess whether performance differences between models were statistically significant.

### Data and materials

The prediction scores from the Random Forest classifier (full model) are available on FigShare: https://doi.org/10.6084/m9.figshare.31058182

All the code is available in GitHub: https://github.com/cancergenetics/context_specific_paralog_SL

## Authors’ contributions

Conceptualization: CJR, NK

Data curation: NK, HA

Formal analysis: NK, HA

Funding Acquisition: CJR

Methodology: NK, HA

Software: NK, HA

Supervision: CJR, DA

Visualization: NK

Writing - original draft: NK, CJR

Writing – review & editing: NK, HA, DJA, CJR

## Acknowledgements

We thank Prashant Gupta and members of the Ryan lab for critical reading of the manuscript.

## Supplementary information

**Figure S1.**
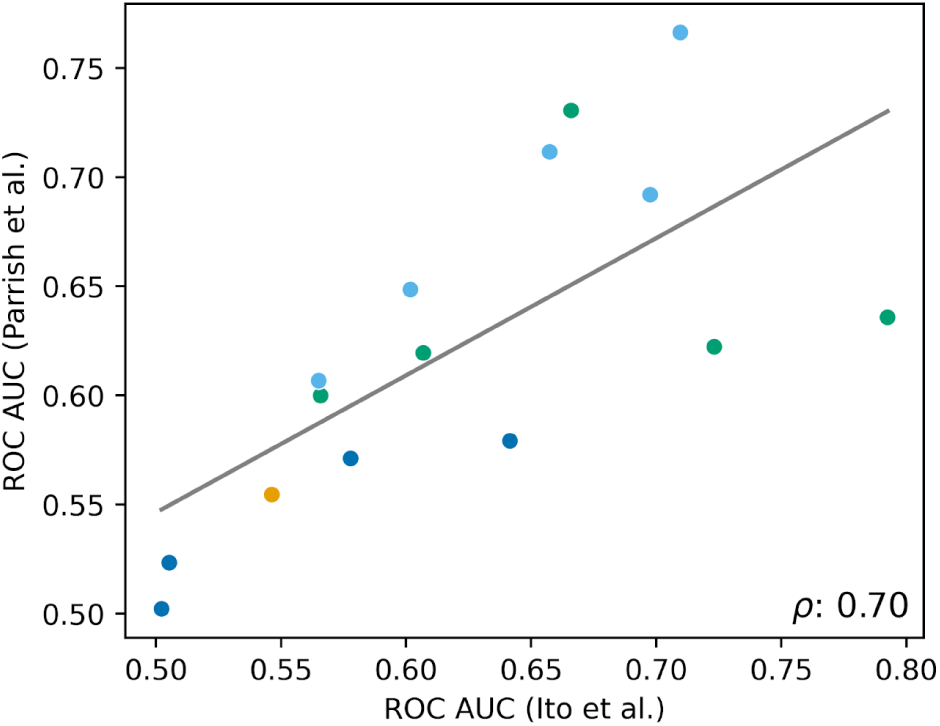
Consistency of feature-level predictive performance across two independent combinatorial CRISPR screens. Each point represents a single predictive feature, plotted by its ROC AUC in the Ito et al. dataset (x-axis) and its ROC AUC in the Parrish et al. dataset (y-axis). Synthetic lethality labels were harmonized across both datasets by reprocessing [15] using the Gemini genetic interaction scoring pipeline [51]. A strong correlation was observed between the two datasets, indicating consistent feature performance: Pearson’s ρ = 0.70 (p = 0.0038) and Spearman’s r = 0.79 (p = 0.00047). This agreement suggests that key features such as gene essentiality, GO-based activity scores, and PPI-derived metrics retain predictive value across distinct experimental conditions.

**Figure S2.**
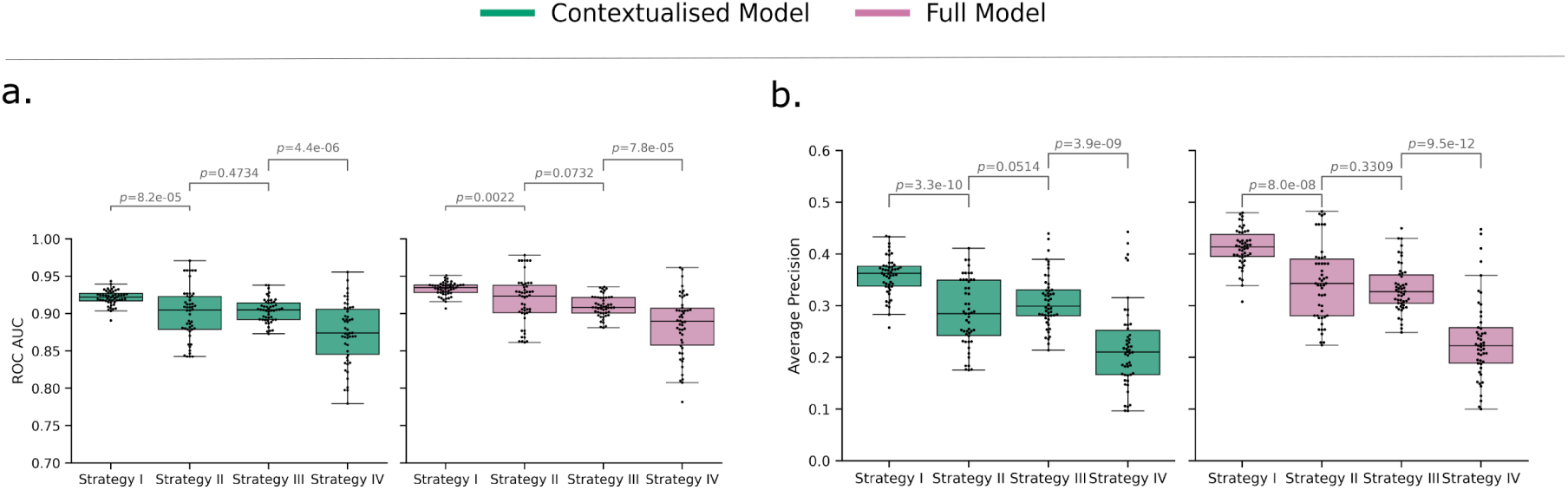
Performance comparison of context-specific classifiers across four validation schemes. Box plots show the distribution of ROC AUC (a) and average precision (b) scores across 50 runs (k=5, n=10 cross validation) for each individual setting: Strategy I (random partitioning), Strategy II (unseen cell lines), Strategy III (unseen gene pairs), and Strategy IV (unseen gene pairs and unseen cell lines) for both the Contextualised and Full Models. Statistical comparisons were performed using t-tests.

**Figure S3.**
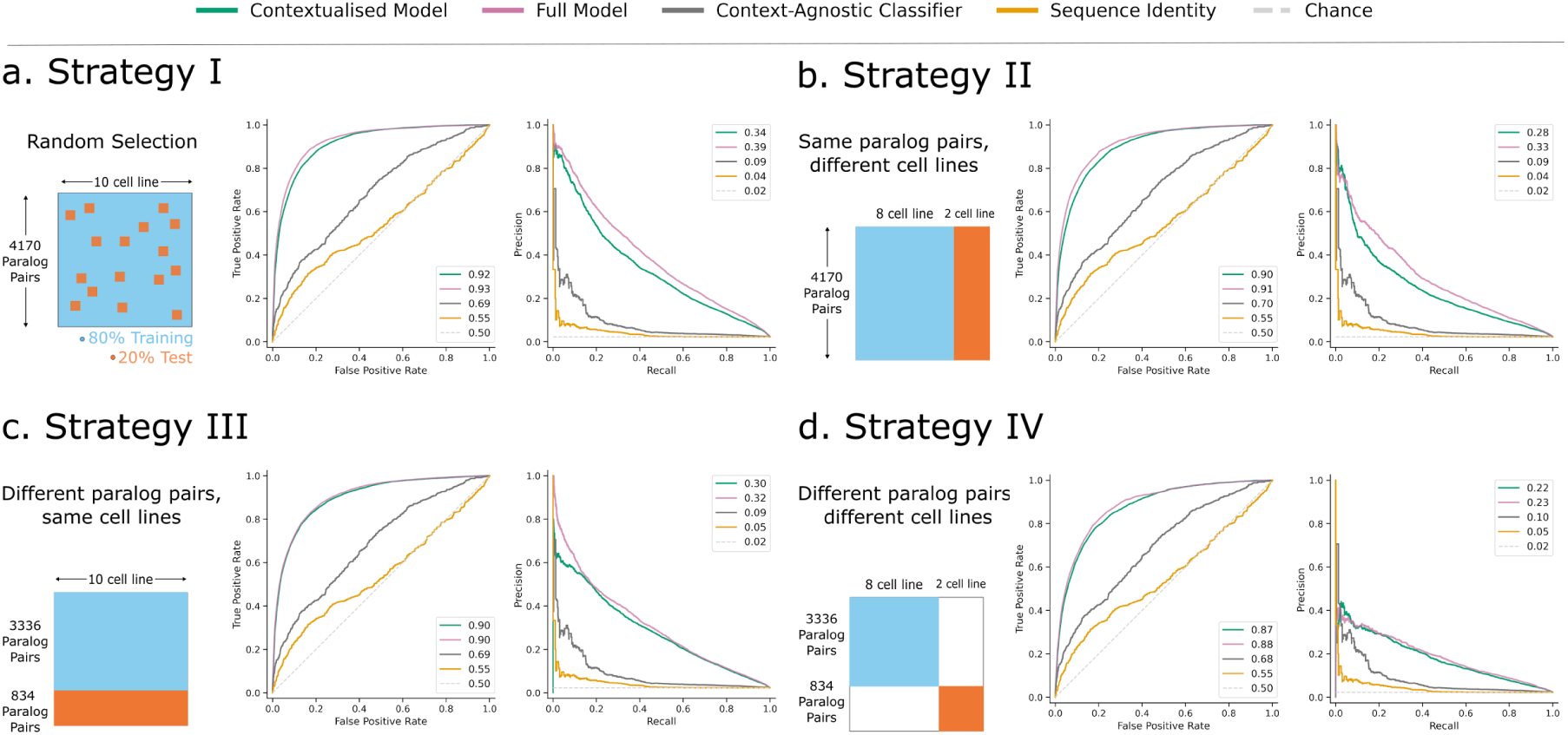
Performance evaluation of the context-specific random forest classifiers across four validation schemes after artificially reducing the data in the training set. (a-d) ROC and Precision-Recall (PR) curves visualize the performance of the context-specific classifier (contextualised and full models) on reduced training data, which allows comparison of Strategies I, II, and III with Strategy IV. The dashed line indicates random performance (chance).

**Figure S4.**
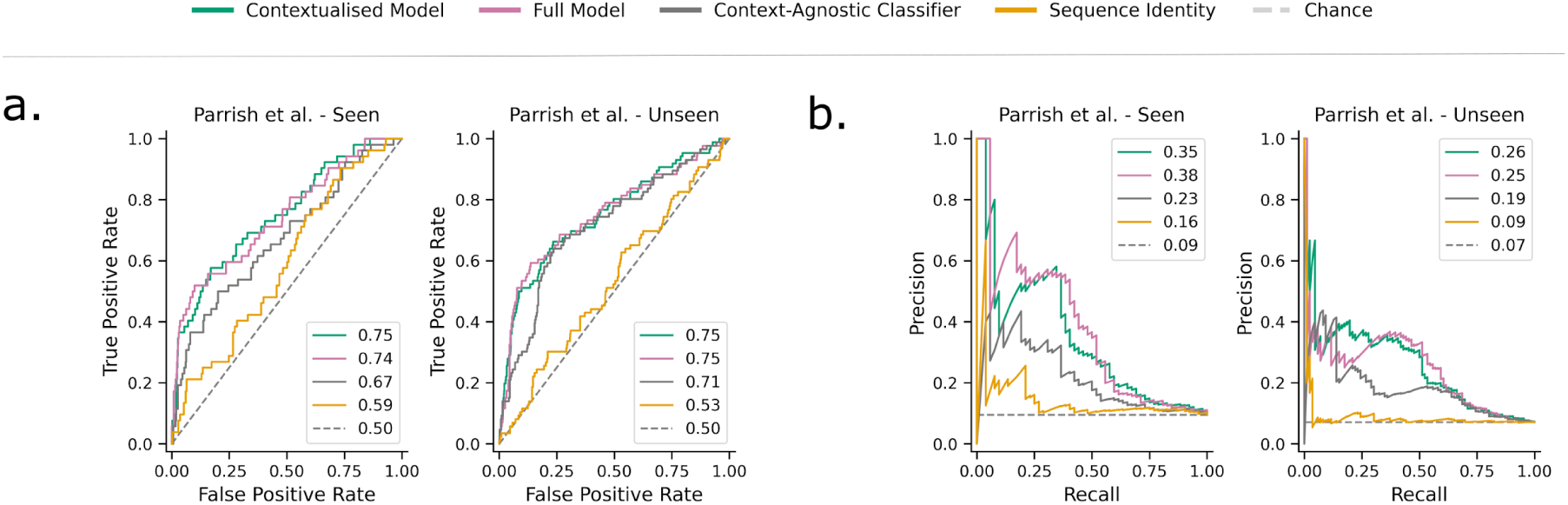
Performance of the context-specific classifier on the Parrish et al. dataset. (a) ROC and (b) precision-recall curves comparing the predictive performance of contextualised and full model classifiers on both seen and unseen subsets originated from the Parrish et al. dataset reprocessed with Gemini.

